# Neural regulation of hatching in amphibians

**DOI:** 10.64898/2026.11.09.765040

**Authors:** María José Salazar-Nicholls, Corey A. H. Allard, Astrid K. Lisondro-Arosemena, Olivia D. Leto, Stephanie Krueger, Nicholas W. Bellono, Karen M. Warkentin

## Abstract

Many animal embryos hatch in response to threats or opportunities, yet how cue sensing activates hatching is unknown. We screened red-eyed treefrog (*Agalychnis callidryas*) eggs in the field with small molecules to discover that the neurotransmitter acetylcholine is a key mediator of hatching enzyme release. Using behavioral pharmacology, microscopy, and transcriptional profiling we identify direct cholinergic input on hatching gland cells that triggers enzyme release for egg rupture, and serotonergic transduction of embryonic hypoxia sensing. Independent hypoxia-sensing and mechanosensory predation-detection pathways converge on a hatching decision that elicits acetylcholine-dependent enzyme release and coordinated hatching-specific behavior. Cholinergic regulation of hatching enzyme release is conserved in a distantly related frog and salamander, suggesting a shared ancestral hatching mechanism. These results provide a neural basis for hatching plasticity which may have contributed to the evolution of diverse hatching modes in amphibians.

## INTRODUCTION

Hatching is a crucial life history transition for most animals, and many have the ability to adaptively hatch in response to environmental cues, within a period of developmental competence^1^. Among terrestrial vertebrates, amphibians have evolved an extraordinary diversity in reproductive modes and adjust hatching timing in response to many stimuli including hypoxia, light, heat, dehydration, and chemical and physical cues from predators, pathogens, and parents^2–10^. Some can delay hatching for months, waiting for conditions that enable larval survival, while others can hatch within seconds to escape an attacking predator^2, 11^. In most amphibians, enzymatic digestion of the egg capsule is crucial to enable embryo exit^12, 13^, but what regulates hatching enzyme release by hatching gland cells is unknown. Moreover, since diverse environmental cues lead to hatching^13, 14^, what links their sensation to hatching enzyme release?

To address this question, we leveraged the specialized hatching mode of red-eyed treefrogs (*Agalychnis callidryas*) to investigate the molecular and cellular mechanisms of hatching plasticity. The arboreal embryos of these frogs can spend a week in their egg capsule if undisturbed, but hatch earlier and very rapidly in response to threats (**Figure 1A, 1B**)^15, 16^. As soon as hatching gland cells become functional, at embryonic age 3 days, hypoxia cues hatching from flooded eggs^17–19^. At 4 days, development of the lateral line and vestibular mechanosensory systems enables escape from predators such as egg-eating snakes and wasps^18, 20, 21^. Hatching involves stereotyped hatching-specific behavior and acute release of hatching enzyme to rapidly rupture the egg capsule (**Figure 1B**)^12, 22^. Accordingly, these embryos develop an extraordinarily large number of hatching gland cells and accumulate large hatching enzyme reserves^19^ (**Figure 1C–E**). How embryonic amphibians sense hypoxia and how the decision to hatch is transduced to hatching gland cells is unknown. Our investigation revealed serotonin-mediated hypoxia sensing and a conserved cholinergic mechanism that controls hatching enzyme release across distantly related amphibians.

**Fig. 1.**
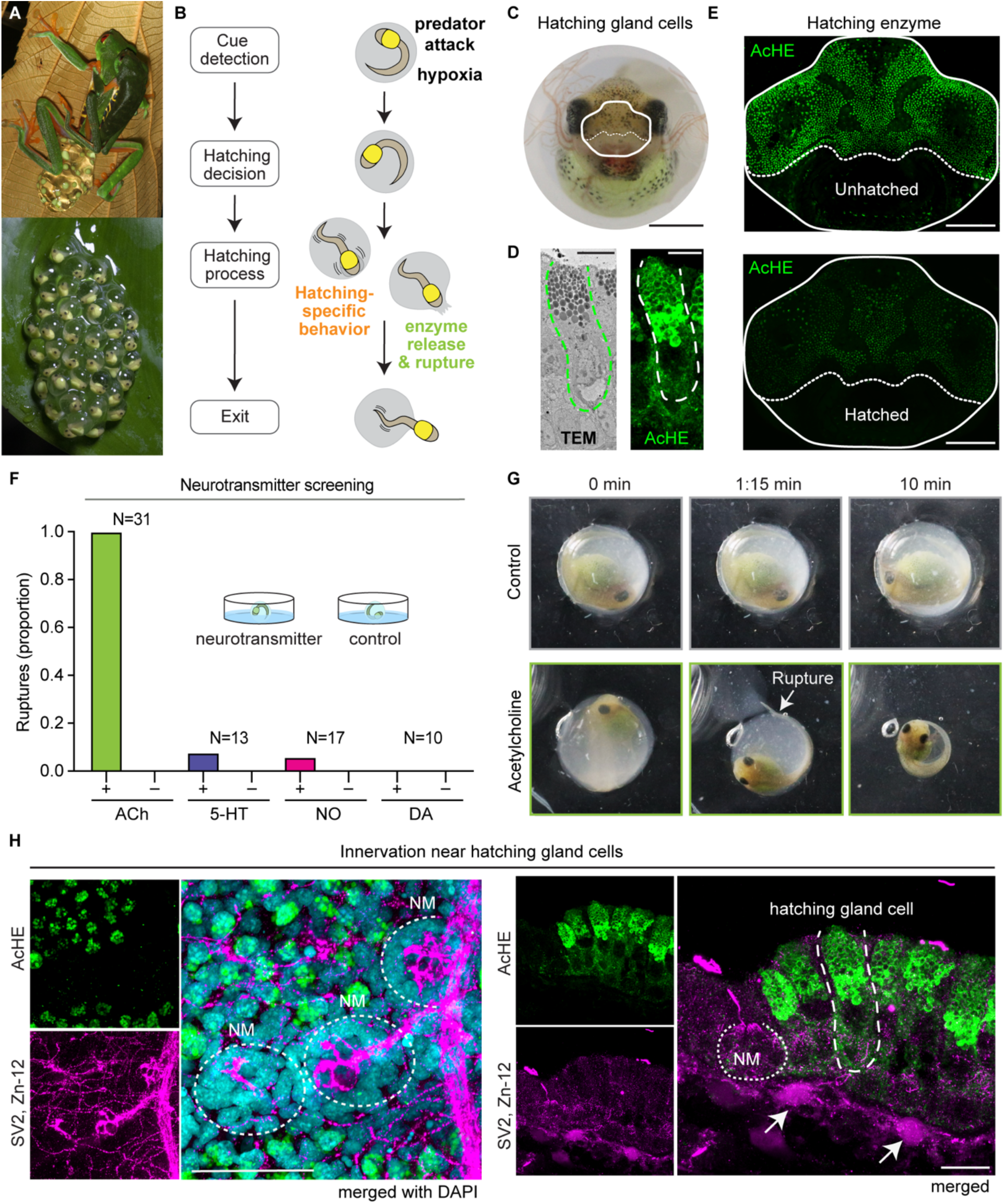
Acetylcholine triggers hatching enzyme release in red-eyed treefrogs. (A) *Agalychnis callidryas* lay arboreal eggs (top) that can hatch after 3 days of development (bottom). (B) Threat cues stimulate embryos to escape by a rapid hatching process. (C) Frontal view of an embryo *in ovo* indicating snout area (white line); hatching gland cells are concentrated on the dorsal snout (above dotted white line). Scale is 1 mm. (D) Section through hatching gland cells: TEM image (left) and fluorescent image with AcHE antibody (right), with dashed outlines around individual HGC; scale bars are 10 μm. (E) Fluorescent frontal images showing hatching enzyme reserves marked with AcHE antibody (green) before and after hatching in 5-day-old embryos; scale bars are 200 μm. (F) Neurotransmitter screening—proportion of egg membrane ruptures during 10 min exposure to neurotransmitters, with paired sibling controls: acetylcholine (ACh, 1 mM), serotonin (5-HT, 500 μm), norepinephrine (NO, 1 mM), dopamine (DA, 1 mM). Samples sizes are indicated; inset shows incubation set-up. (G) Video frames from recording of a 4-day-old embryo during 10 min acetylcholine exposure, showing initial membrane rupture, and paired sibling control. (H) Frontal view (left) and coronal section (right) of embryo snout skin, showing innervation marked with synaptic vesicle (SV2) and neural surface (Zn-12) antibodies in magenta and hatching glands cells marked with AcHE antibody in green. Left: lateral line nerve, neuromasts (NM), and intricate skin innervation around densely concentrated hatching gland cells. Nuclei are marked with DAPI (cyan) in merged image; scale is 50 μm. Right: Neural cells (arrows) below the basal lamina near hatching gland cells and neuromasts. Scale is 10 μm.

## RESULTS

### Cholinergic signaling mediates hatching enzyme release in red-eyed treefrogs

Considering the close coordination between cue sensing and hatching enzyme release, we hypothesized that the activation of hatching gland cells is controlled by neural signaling. To test this, we treated field-collected 4-day-old *A. callidryas* embryos with four neurotransmitters selected for their roles in glandular activation and hypoxia-cued behavior (acetylcholine^23^, norepinephrine^24^, dopamine^25^, and serotonin^26^) and observed egg membrane integrity and embryo behavior. Strikingly, all embryos exposed to acetylcholine rapidly ruptured their egg membrane, indicating enzymatic digestion (**Figure 1F, 1G**, **S1A, movie S1**). Other neurotransmitters did not induce membrane rupture. Rupture occurred at the embryo’s snout and perivitelline fluid leaked out as soon as the embryo changed position, leading to capsule collapse without hatching-specific behavior (**Figure 1G, S1B and movie S1**). Histological examination of whole and sectioned embryos revealed a potential endogenous source of cholinergic neurotransmission to the hatching gland cells. We found that the snout epithelium, where hatching gland cells are concentrated, is densely innervated (**Figure 1H and S1C-D**) and has cholinergic cells located 20 μm from the hatching gland cells, just below the basal lamina **(Figure 2A**). Although synapses were evident on hair cells of nearby neuromasts, we did not find them on the hatching gland cells (**Figure S2**), suggesting nonsynaptic volume transmission that is often observed in bulk cholinergic signaling systems^27^.

**Fig. 2.**
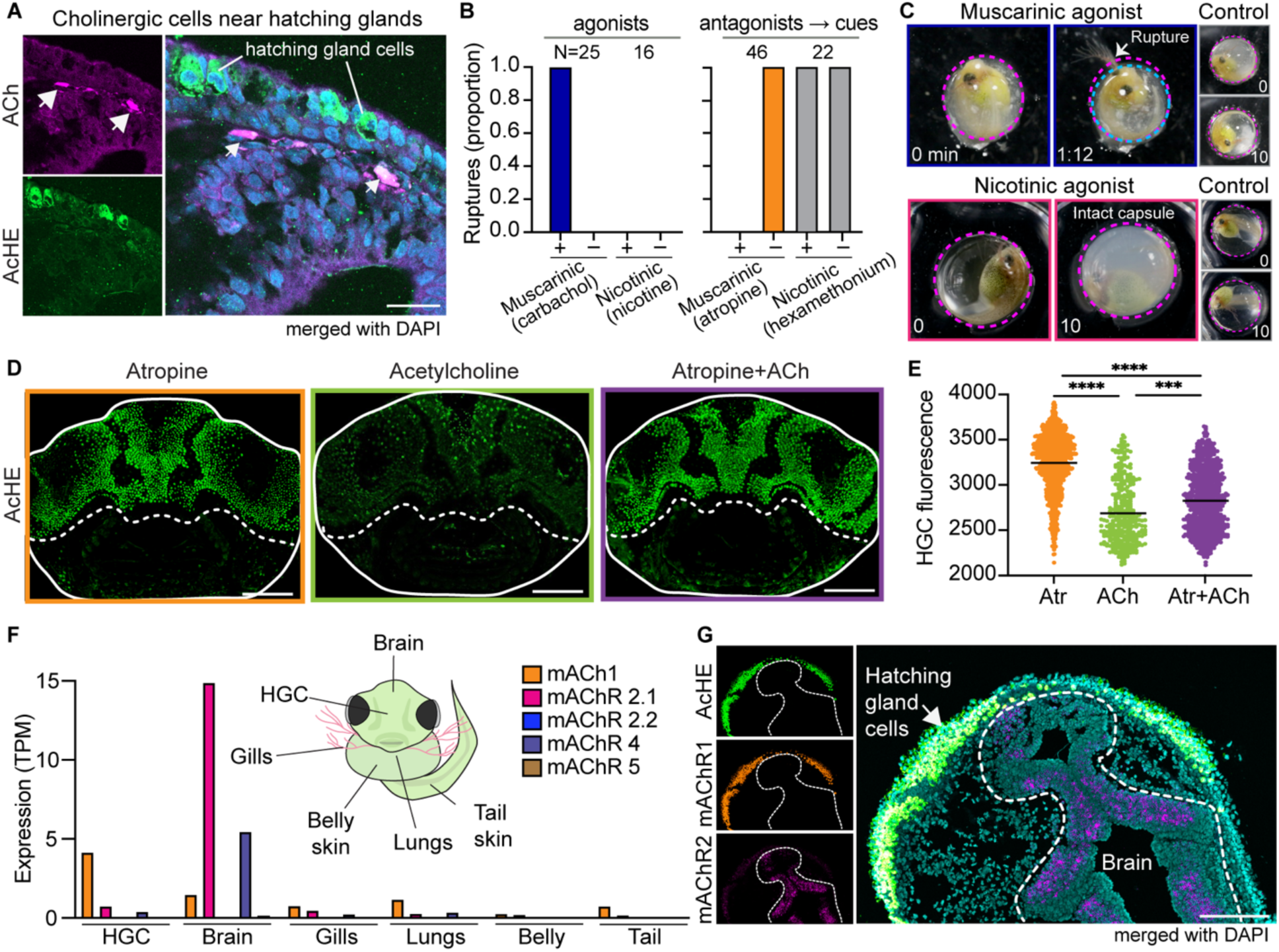
Signaling through muscarinic acetylcholine receptor mAChR1 activates hatching gland cells to release hatching enzyme in red-eyed treefrogs. (A) Section through an embryo’s snout skin showing cholinergic cells (arrows) marked with ACh antibody (magenta) just underneath the basal lamina of the epidermis, below hatching gland cells marked with AcHE antibody (green); scale is 20 μm. (B) Cholinergic receptor type screen showing proportion of egg membrane ruptures during (left) 10 min exposure to agonists and (right) 10 min exposure to antagonists followed by hatching cues (30 min hypoxia or 10 min simulated predation); sample sizes are indicated. The muscarinic agonist carbachol (5 mM) consistently induced membrane rupture; nicotine (500 µM) did not. The muscarinic antagonist atropine (100 μM) consistently blocked membrane rupture and hatching, while all controls and embryos treated with the nicotinic antagonist hexamethonium chloride (500 μM) hatched successfully in response to cues. (C) Video frames from muscarinic and nicotinic receptor agonist screenings of 4-day old embryos observed for egg membrane rupture during 10 min of pharmacological exposure and paired controls; time (min) is indicated. Magenta outlines intact egg membranes; blue outlines ruptured, shrinking membrane (carbachol treatment only, note protruding external gill). (D) Fluorescent frontal images of decapsulated embryos’ snouts showing hatching enzyme marked with AcHE antibody (green) after 10 min exposure to atropine (a muscarinic receptor antagonist), acetylcholine, or atropine followed by acetylcholine; scale bars are 200 μm. Detected cell counts (mean ± SD): atropine 3243 ± 317, acetylcholine 2688 ± 345, atropine + acetylcholine 2815 ± 338. (E) Fluorescence quantification of hatching enzyme in cells detected in frontal images (D); Kruskal-Wallis test P<0.0001 (****) followed by Dunn’s multiple comparisons, Atropine vs. ACh or both P<0.0001, ACh vs. both P=0.001 (***). (F) mRNA expression profile of muscarinic receptors across embryo tissues; inset shows tissues collected for transcriptomic analysis. (G) Coronal section through embryo head, showing localization of hatching enzyme (green) and mAChR1 (orange) mRNA in the hatching gland cells and mAChR2 (magenta) in the brain, visualized by fluorescent in situ hybridization. Nuclei are stained with DAPI (cyan); scale bars are 200 μm.

We next sought to identify the acetylcholine receptors involved in enzyme release. The nicotinic acetylcholine receptor (nAChR) agonist nicotine did not induce membrane rupture, and all embryos exposed to the nAChR antagonist hexamethonium hatched normally in response to cues (**Figure 2B–C and S3A**). In contrast, exposure to the muscarinic acetylcholine receptor (mAChR) agonist carbachol triggered rapid membrane rupture in every tested embryo, and the mAChR antagonist atropine blocked hatching (**Figure 2B–C and S3A, movie S2**). Atropine-treated embryos showed persistent behavioral attempts to hatch in response to both hypoxia and simulated predation (a mechanical egg-jiggling stimulus)^18^, forcefully pushing and shaking their heads against the membrane, but no membrane rupture occurred (**movie S3**). Furthermore, in decapsulated embryos, acetylcholine exposure was sufficient to deplete hatching enzyme reserves, indicating enzyme release, and this depletion was prevented by pre-treatment with atropine (**Figure 2D–E**). Thus, we hypothesized that hatching gland cells themselves could be the target of cholinergic signaling. Tissue-specific RNA sequencing demonstrated that mAChR1 is specifically enriched in the snout epithelium where hatching gland cells are concentrated, in contrast to mAChR2 that was enriched in the brain (**Figure 2F**). Furthermore, RNA-FISH showed transcripts encoding mAChR1 localized to hatching gland cells, while mAChR2 was found in the brain (**Figure 2G**). We conclude that the hatching gland cells release hatching enzyme in response to mAChR1-mediated cholinergic signaling downstream of diverse sensory inputs.

### Serotonergic signaling mediates hypoxia-cued hatching

As enzymatic digestion of the membrane is essential for hatching to occur, we next sought to understand how different sensory inputs converge on hatching enzyme release. Embryonic mechanosensing via lateral line and vestibular systems is known to mediate hatching in response to egg-predator attacks^20, 21^. However, even though hypoxia-cued hatching is widespread in amphibians^2^, embryonic oxygen-sensing mechanisms are unknown. In more developed, larval-stage amphibians and fishes, oxygen sensing involves serotonergic neuroepithelial cells in the gills and 5-HT_3_ receptors expressed in gill neurons^26, 28, 29^. Consistent with these findings, we found embryonic *A. callidryas* gills to be densely innervated, with serotonergic neuroepithelial cells present, and tissue enriched for the serotonin receptor 5-HT_3A_ (**Figure 3A–C**). Furthermore, serotonin-treated embryos flooded with hypoxic water hatched more rapidly than their unexposed siblings (**Figure 3D and F, movie S4**) and the 5-HT_3_ receptor antagonist MDL72222 blocked or severely delayed embryos’ hatching response to hypoxia (**Figure 3E, 3G, movie S4**). These results support serotonergic signaling received by 5-HT_3_R as a key mechanism in hypoxia-cued hatching, elucidating a distinct link between sensory input and the hatching response.

**Fig. 3.**
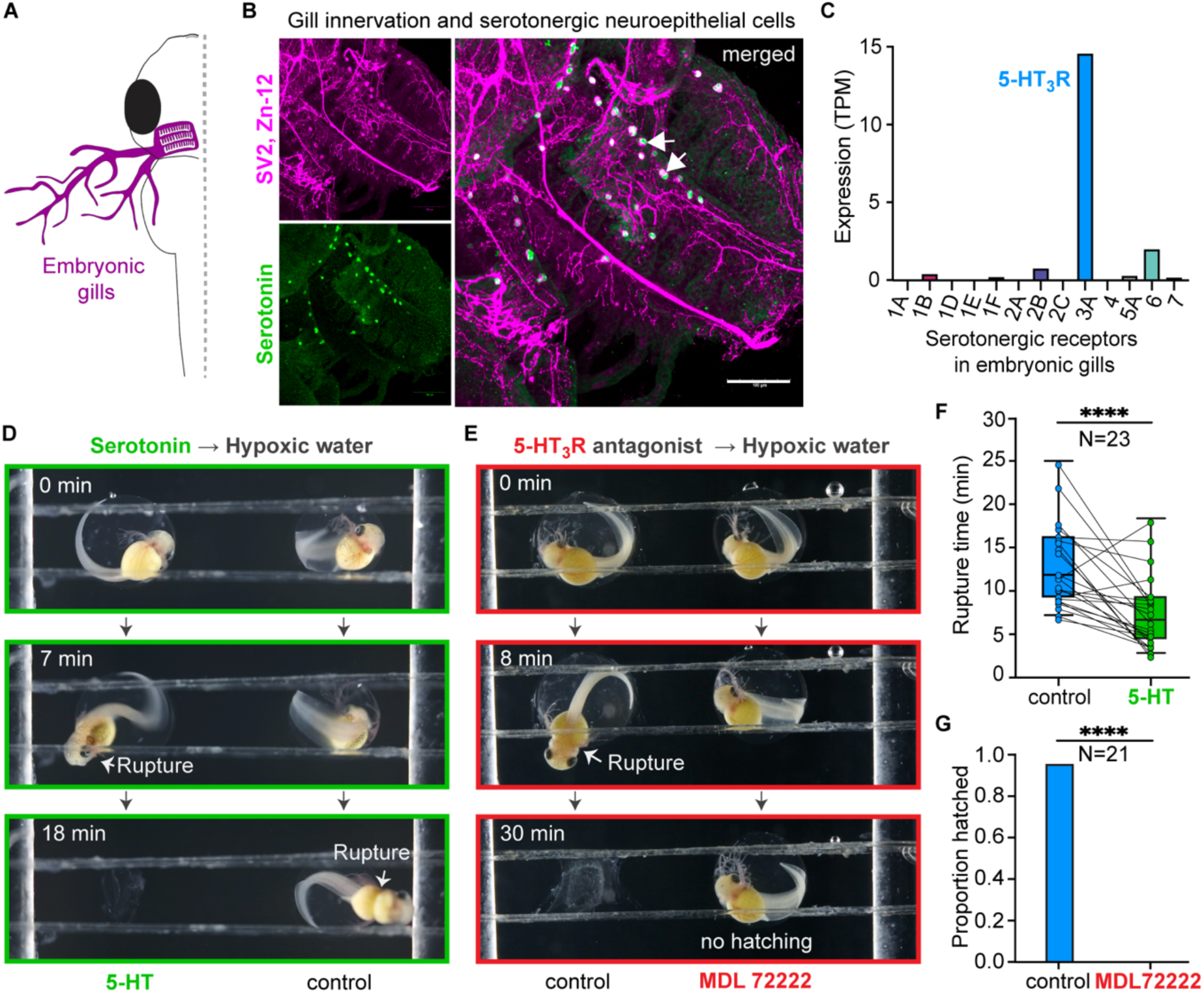
Serotonergic neuroepithelial cells and hypoxia sensing in red-eyed treefrog embryos. (A) Diagram of embryonic gills in A*galychnis callidryas*. (B) Embryo’s internal gills showing innervation marked with synaptic vesicle and neural surface antibodies (SV2 and Zn-12, magenta) and serotonergic cells marked with 5-HT antibody (green). Merged image shows serotonergic neuroepithelial cells (arrows); scale is 100 μm. (C) Transcript abundance of serotonin receptor subtypes in embryonic gills. TPM = transcripts per million. (D) Video frames from recording of sibling embryos in hypoxic water, showing accelerated hatching of serotonin pre-exposed (500 µM, 10 min) individual compared to control; times are indicated. (E) Video frames showing no hatching response to hypoxia by an embryo pre-exposed to the 5-HT_3_ receptor antagonist MDL 72222 (500 µM, 10 min) and hatching of its control sibling. (F) Latency of hypoxia-cued hatching response for embryos pre-exposed to serotonin and paired sibling controls, measured as time from flooding to membrane rupture. Data represent individual embryos with lines connecting pairs; box plots show median and interquartile range (IQR), whiskers show 5th and 95th percentiles, N represents paired trials. Wilcoxon matched pairs signed rank test P<0.0001 (****). (G) Proportion of embryos pre-exposed to 5-HT_3_R antagonist MDL 72222 and paired sibling controls that hatched in response to hypoxia, Fisher’s exact test P<0.0001.

### Sensory signals converge to elicit coordinated enzyme release and capsule exit

In red-eyed treefrogs, the onset of oxygen sensing, hatching ability, lateral line function, and vestibular mechanosensing are separated, sequential developmental events^18–20, 30^. To probe the independence of sensory and effector mechanisms in cued-hatching pathways, we blocked oxygen-sensing with the 5-HT_3_R antagonist MDL72222 and asked if embryos still hatched in response to a mechanosensory hatching stimulus (egg-jiggling) or released hatching enzyme upon exposure to acetylcholine. All jiggled embryos hatched, and all acetylcholine-exposed embryos rapidly ruptured their membrane (**Figure 4A, S3B**). Thus, blocking 5-HT_3_R did not affect embryos’ ability to detect a mechanosensory cue, release hatching enzyme to rupture the membrane, or successfully exit the egg; it only impaired the cue-detection step in the hypoxia-cued hatching pathway (**Figure 4A**). Similarly, mAChR-antagonist exposure did not alter embryos’ oxygen-sensing, mechanosensing, or hatching-specific behavior; only hatching enzyme release was impaired, blocking membrane rupture (**Figure 4A**). Moreover, neither hatching enzyme release nor hatching-specific behavior independently enabled hatching; exit from the capsule only occurred when embryos performed both in a coordinated manner (**Figure 4B**). Together, our results suggest that multiple independent sensory pathways converge at a hatching decision that initiates both hatching-specific behavior and cholinergic signaling to release hatching enzyme for egg rupture, which together enable rapid exit (**Figure 4C**).

**Fig. 4.**
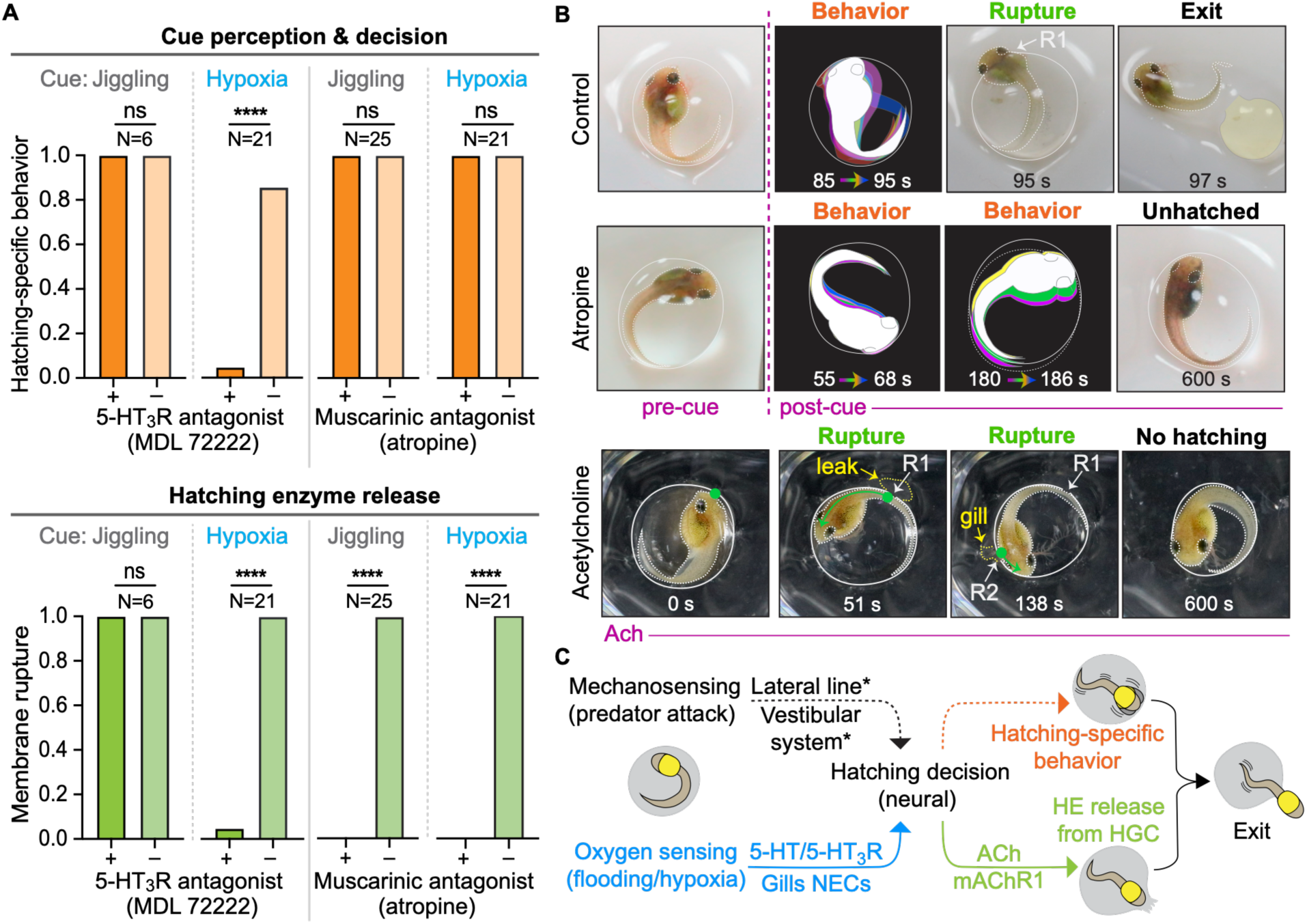
Independent neural modulation of cue sensing and hatching enzyme release. (A) Occurrence of (top) hatching behavior, an indicator of cue perception and the decision to hatch, and (bottom) egg membrane rupture, an indicator of hatching enzyme release, after exposure to a 5-HT_3_ receptor antagonist (MDL 72222) or muscarinic receptor antagonist (atropine), followed by either a mechanosensory or hypoxia cue, with paired controls. Fisher’s exact test P<0.0001 (****). (B) Cued control embryo (top) performs hatching-specific behavior and releases enzyme to rupture membrane (R1), then exits immediately. Its sibling pre-exposed to atropine (middle: 100 μM, 10 min) performs hatching-specific behavior repeatedly but cannot release enzyme to rupture capsule, so remains unhatched. Acetylcholine exposure (1 mM, bottom) induces rapid enzyme release; ruptures (R1, R2) are evident (fluid leak, gill protrusion) once embryo moves (green dots and arrows), but embryo performs no hatching-specific behavior and does not exit. Frame times are from onset of mechanosensory cue (control, atropine) or ACh treatment. Kymographs show embryo positions in frames across periods of hatching-specific behavior. (C) Model of multi-component cued-hatching pathway, with distinct cues and sensors providing input to a hypothesized CNS center that coordinates motor and glandular output for successful exit from the _egg. *20, 21_

### Cholinergic signaling mediates hatching enzyme release in amphibians

Although red-eyed treefrogs release hatching enzyme acutely to hatch in seconds, other amphibians release hatching enzyme gradually and may take hours or days to hatch^12^. Is cholinergic signaling a specific adaption for rapid cued hatching, or could it represent a conserved component of hatching machinery more broadly? To assess the potential generality of acetylcholine-mediated hatching enzyme release, we tested embryos of a distantly related frog, a salamander, and a fish (**Figure 5A**). In the African clawed frog *Xenopus tropicalis* and axolotl *Ambystoma mexicanum*, acetylcholine exposure triggered hatching enzyme release, but in the zebrafish *Danio rerio* it did not (**Figure 5A and S4**). Moreover, transcripts encoding hatching enzyme and mAChR1 colocalized within individual hatching gland cells in the head epithelium of both *X. tropicalis* and *A. mexicanum* (**Figure 5B**), but we did not detect mAChR1 transcripts localized in the hatching gland cells of zebrafish (**Figure 5B**). Our results suggest that the mAChR1-mediated cholinergic control of HE release is likely ancestral and broadly shared in amphibians (**Figure 5C**).

**Fig. 5.**
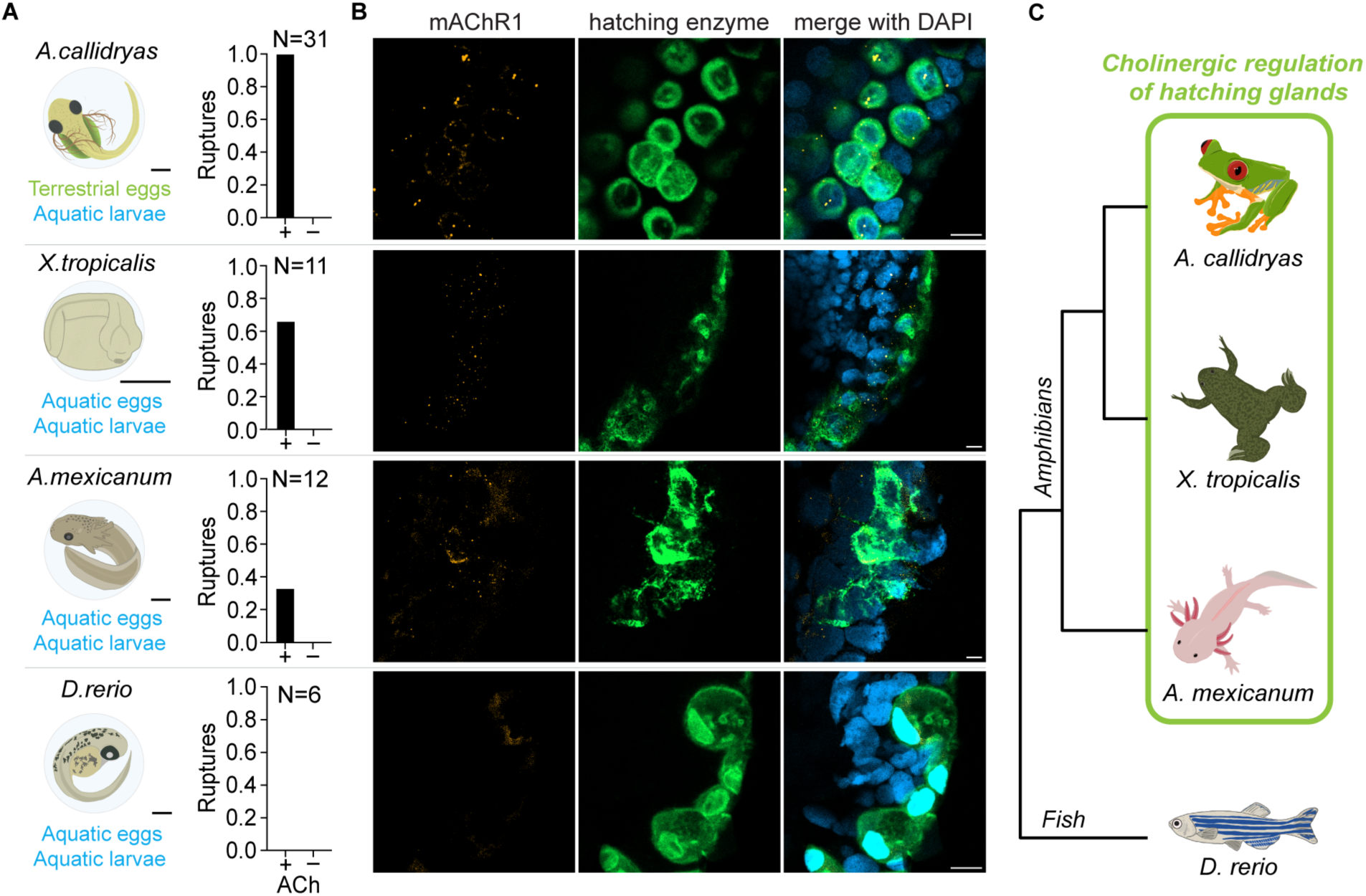
Cholinergic regulation of hatching enzyme release in amphibians. (A) Left: Illustrations of hatching-competent embryos of each tested species, with egg and larval environments listed. Scale bars are 1 mm. Right: Incidence of egg capsule rupture by embryos of each species in response to acetylcholine exposure (1 mM; times: *A. callidryas* and *X. tropicalis* 10 min, *A. mexicanum* 45 min, *D. rerio* 40 min), with paired controls. (B) Hatching gland cells of each species showing mRNA transcripts of hatching enzyme (green) and mAChR1 (orange, in amphibians) as visualized by fluorescent in situ hybridization. Nuclei were stained with DAPI (cyan); scale bars 10 μm. (C) Phylogenetic relationship of tested species, showing cholinergic regulation of hatching gland cells in amphibians.

## DISCUSSION

### Regulation of hatching enzyme release in amphibians

We leveraged the best-studied example of cued hatching in amphibians, the red-eyed treefrog, to discover neural mechanisms underlying escape hatching. Cholinergic signaling to hatching gland cells elicits hatching enzyme release, forming a mechanism shared among the amphibian species studied here. Considering this mechanism is present in two distantly related frogs and a salamander that differ dramatically in reproductive modes, embryonic adaptations, and hatching modes and diverged about 270 mya^31^, cholinergic control of hatching enzyme release is likely ancestral to Batrachia, the clade that includes frogs and salamanders. It may be shared more broadly across amphibians, but the mechanism regulating hatching enzyme release in the third amphibian clade, caecilians, is unknown.

Amphibians can hatch at developmental stages from tailbud to tiny frog^32, 33^, and embryos’ sensory abilities vary accordingly. At hatching, *Xenopus* have a light-sensing pineal gland and a few mechanosensory neuromasts^34, 35^, while red-eyed treefrogs have well developed eyes and ears, an extensive lateral line, and oxygen and ammonia chemosensing^20, 21, 36^. Cued hatching has not been documented in *X. tropicalis* or *A. mexicanum.* Both hatch more slowly than *A. callidryas*, with far fewer hatching gland cells and no specialized hatching-specific behavior, as may be common among aquatic amphibian embryos^37, 38^. In *A. callidryas*, complex cue assessment and multisensory integration for hatching decisions^10, 21, 39, 40^ and coordination of hatching-specific behavior with hatching enzyme release indicate neural control of hatching, and the cholinergic neurons near hatching gland cells provide a mechanism. However, for amphibians such as *Xenopus* and toads (*Anaxyrus*) that hatch with less developed nervous systems and simpler or no regulation by environmental cues^33, 37^, non-neuronal cholinergic signaling might play a role in HE release. Consistent with this, mAChR1 is commonly implicated in non-neuronal cholinergic signaling, suggesting it could function in both systems. Thus, cholinergic regulation of hatching enzyme release may be simply developmentally gated or linked to upstream regulatory pathways that vary in complexity.

### Divergent regulation of hatching gland function in amphibians and fishes

Hatching enzyme was present in the common ancestor of bony fishes and is orthologous across tetrapods and ray-finned fishes^13^. However, the gland cells that produce it originate from mesoendoderm during gastrulation in teleosts and from ectoderm during neurulation in frogs^41^. We found that the molecular signal for hatching enzyme release has also diverged. In teleost fishes, neurohormonal signaling is hypothesized to activate hatching enzyme release^25, 42^ and two distinct pathways have been identified. In halibut, projections from a cluster of light-sensitive hindbrain neurons innervate the hatching gland cells^43^. In zebrafish and medaka, hypothalamic neurons release thyrotropin releasing hormone (Trh) into blood vessels that irrigate the hatching gland cells, where a Trh receptor regulates hatching enzyme release^44^. The cholinergic pathway in amphibians is distinct from both identified teleost pathways. Amphibians exhibit remarkable diversity in reproductive ecology and embryonic adaptations^14, 45^. Their diversity in hatching modes^19^ may reflect innovation from a shared core mechanism of hatching enzyme release, opening new possibilities for comparative research in evolution and neurobiology.

### Ecologically grounded embryo neurobiology in the field

Investigation of *A. callidryas’* specialized mode of hatching enabled the discovery of both shared fundamentals of amphibian hatching and embryonic sensory mechanisms that mediate adaptive embryo behavior. This species produces large eggs, with convenient developmental timing and high visibility *in ovo*, throughout an extended annual breeding season. The separation of egg and larval habitats imposes strong selection for cued hatching and facilitates ecological and behavioral research^16, 46, 47^. We expect *A. callidryas*, with their foundational research in embryo cognitive ecology^48^ and growing behavioral^49^ and genetic tools, will provide further insight into mechanisms underlying adaptive embryo behavior and the hatching process. Moreover, our multifaceted methods, designed for feasibility under Neotropical field conditions, offer a tractable approach for integrative experimental research on diverse non-model frog embryos.

## Resource availability

### Lead contact

Further information and requests for resources should be directed to and will be fulfilled by the lead contact, Karen Warkentin.

## Materials availability

This study did not generate new unique reagents.

## Data and code availability

Tissue RNA-seq raw data are deposited at NCBI Sequence Read Archive (SRA) BioProject ID PRJNA1541026 under the accession numbers: SRR41071719 (RNAseq of *A.callidryas* gills), SRR41071720 (RNAseq of *A.callidryas* brain), SRR41071717 (RNAseq of *A.callidryas* tail skin), SRR41071721 (RNAseq of *A.callidryas* HGCs), SRR41071718 (RNAseq of *A.callidryas* yolk skin), SRR41071716 (RNAseq of *A.callidryas* lungs).

Additional data reported in this paper is available through Figshare at https://figshare.com/s/bf1a4cc07a96f620b9db and will be published upon manuscript acceptance.

This paper does not report any original code.

## Acknowledgements

We thank Sally Seraphin, Jessica Whited, and the MCB zebrafish facility at Harvard for gifts of embryos for comparative work. We thank Team Treefrog 2021–2023, particularly Caroline Guevara-Molina, for assistance in Panama and the Gamboa Frog Group, especially Ryan Taylor and Kim Hunter, for their multidisciplinary perspectives, discussions, and feedback during field seasons. We thank Brittany Walsh and the members of the Bellono lab at Harvard University for their logistical and lab assistance, as well as helpful advice and feedback. We thank Jorge Ceballos, Isis Ochoa, Felix Rodriguez, Raineldo Urriola, Rachel Page, and Greg Cohen for scientific and logistical support at STRI and Maria Ericsson for assistance with TEM at Harvard. Caroline Guevara-Molina and Andres Romero-Carvajal provided helpful feedback and valuable discussions that improved this study. We thank Michael Ryan for comments that improved the manuscript.

## Author contributions

Conceptualization, MJSN, CAHA, NWB, and KMW; methodology, MJSN, CAHA, NWB, and KMW; Investigation, MJSN, CAHA, AKLA, ODL, SK, and KMW; validation, MJSN, CAHA; visualization, MJSN, CAHA; data analysis, MJSN, CAHA; writing – original draft, MJSN and KMW; writing – review & editing, MJSN, CAHA, NWB, and KMW; project administration, NWB and KMW; funding acquisition, MJSN, CAHA, AKLA, NWB, and KMW; resources, NWB and KMW; supervision, NWB and KMW.

## Supplemental Information

Document S1. Figures S1–S4 and movie captions Movies S1–S4

## SUPPLEMENTAL FIGURES AND CAPTIONS

**Fig. S1.**
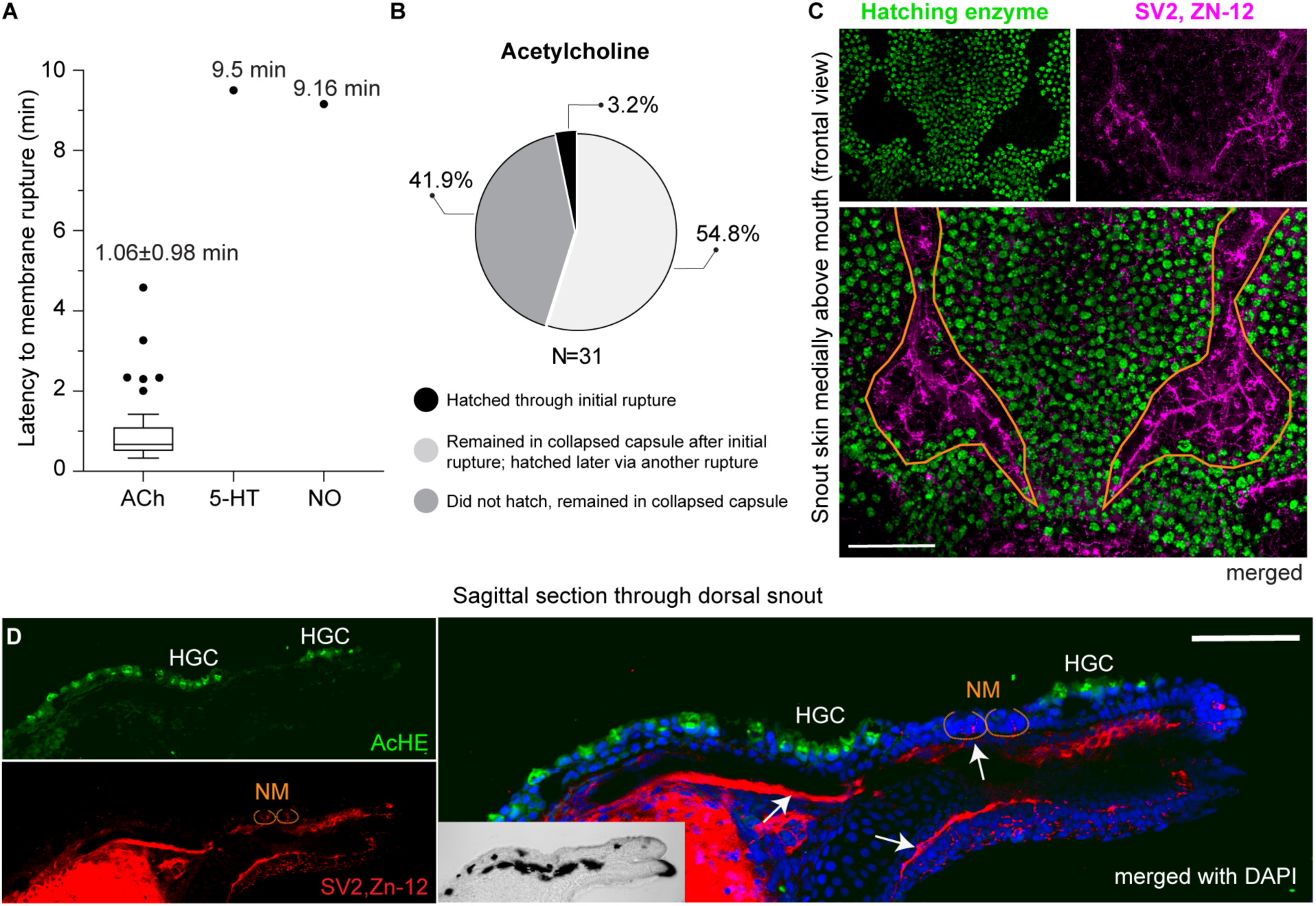
Membrane rupture timing during neurotransmitter screen and innervation near hatching gland cells. (A) Latency to membrane rupture after onset of neurotransmitter exposure in *Agalychnis callidryas* embryos. Data represent individual embryos; for acetylcholine (ACh 1 mM, N=31) box plot shows mean and interquartile range, text shows mean ± SD, whiskers show 5th and 95th percentiles. Serotonin (5-HT 500 μM) and norepinephrine (NO 1 mM) data points indicate times of the single ruptures that occurred in each treatment; no rupture occurred in controls (not plotted). (B) Embryo outcomes during 10 min acetylcholine exposure (1 mM). (C, D) Innervation marked with synaptic vesicle SV2 and neural surface Zn-12 antibodies (magenta or red) near hatching gland cells marked with AcHE antibody (green). (C) Frontal view of a 4-day-old embryo’s medial snout skin showing lateral line (orange dotted lines) and innervation of an area rich in hatching gland cells. (D) Sagittal section of an embryo’s snout (light image inset) showing nerve fibers (red) below the basal lamina, underneath the hatching gland cells (HGC) and neuromasts (NM, dotted orange line). Cell nuclei are marked with DAPI (blue). Arrows show nerve fibers and innervation of neuromasts crossing the basal lamina. Scale bars 50 μm.

**Fig. S2.**
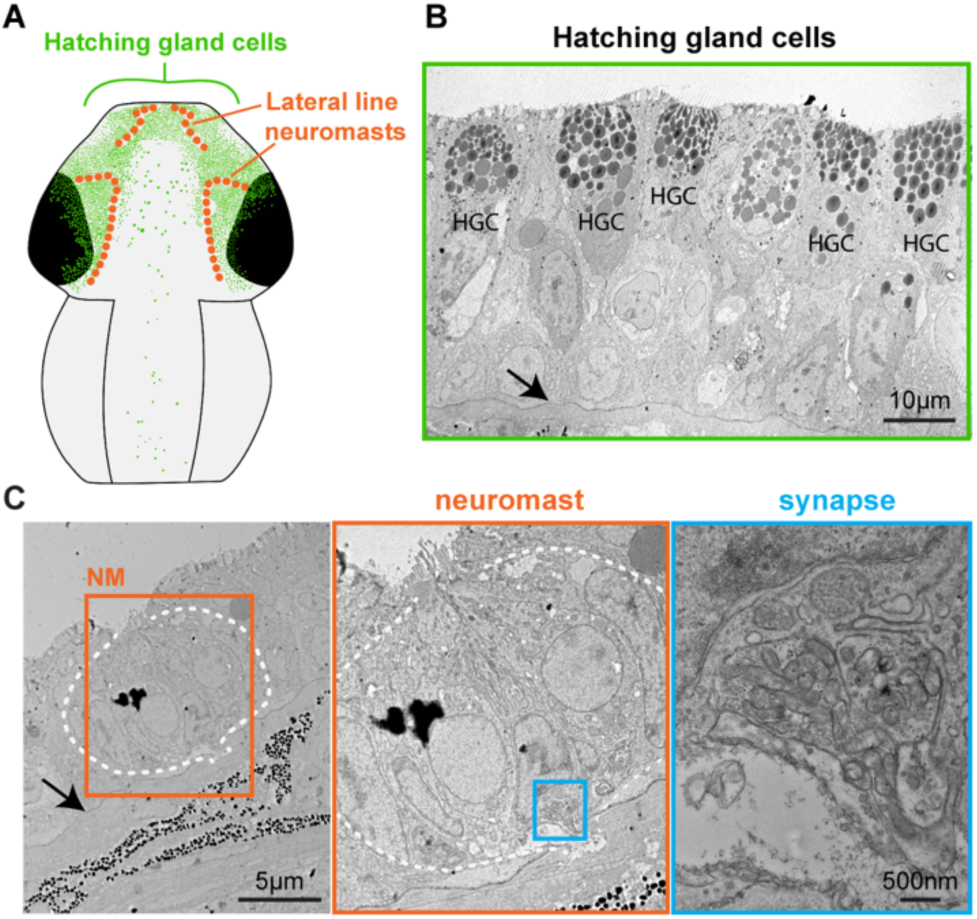
TEM images of hatching gland cells and neuromasts. (A) Diagram of a dorsal view of an *A. callidryas* embryo showing hatching gland cells (green) and lateral line neuromast (orange). (B) Cross-section showing hatching gland cells. Arrow indicates the basal lamina. (C) Cross-section showing a neuromast (white dotted line). Arrow indicates the basal lamina. Insets on the right are enlargements of the boxed area in the left images showing synapse nerve junction crossing the basal lamina to the neuromast (NM). Black arrows indicate basal lamina.

**Fig. S3.**
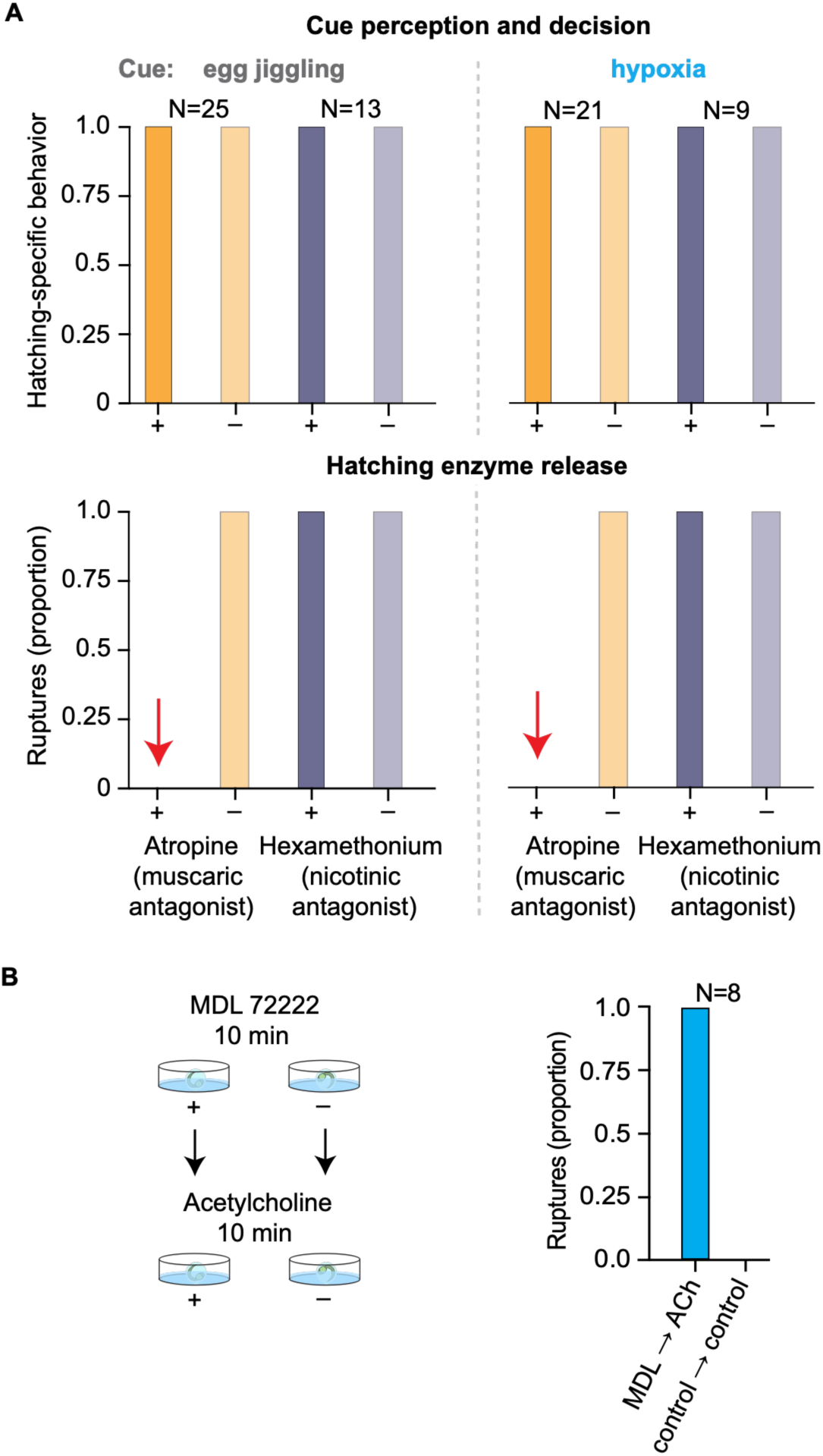
Effect of cholinergic and serotonergic antagonists on cued hatching. (A) Occurrence of (top) hatching-specific behavior, an indicator of cue perception and the decision to hatch, and (bottom) egg membrane rupture, an indicator of hatching enzyme release, after exposure to a muscarinic receptor antagonist (atropine, 1 mM) or nicotinic receptor antagonist (hexamethonium chloride, 500 μM) for 10 min followed by a mechanosensory or hypoxia cue, with paired controls. Red arrows indicate no embryo pre-treated with atropine ruptured its capsule when stimulated with either a mechanosensory or hypoxia hatching cue. (B) Left: diagram shows incubation set-up for sequential pharmacological exposure. Right: Occurrence of egg membrane rupture, an indicator of hatching enzyme release, in embryos exposed first to MDL 72222 (5-HT_3_R antagonist, 500 µM) or a control solution (during which no ruptures occurred) and then to acetylcholine (1 mM) or a control, each for 10 min. Treatment (+), control (–).

**Fig. S4.**
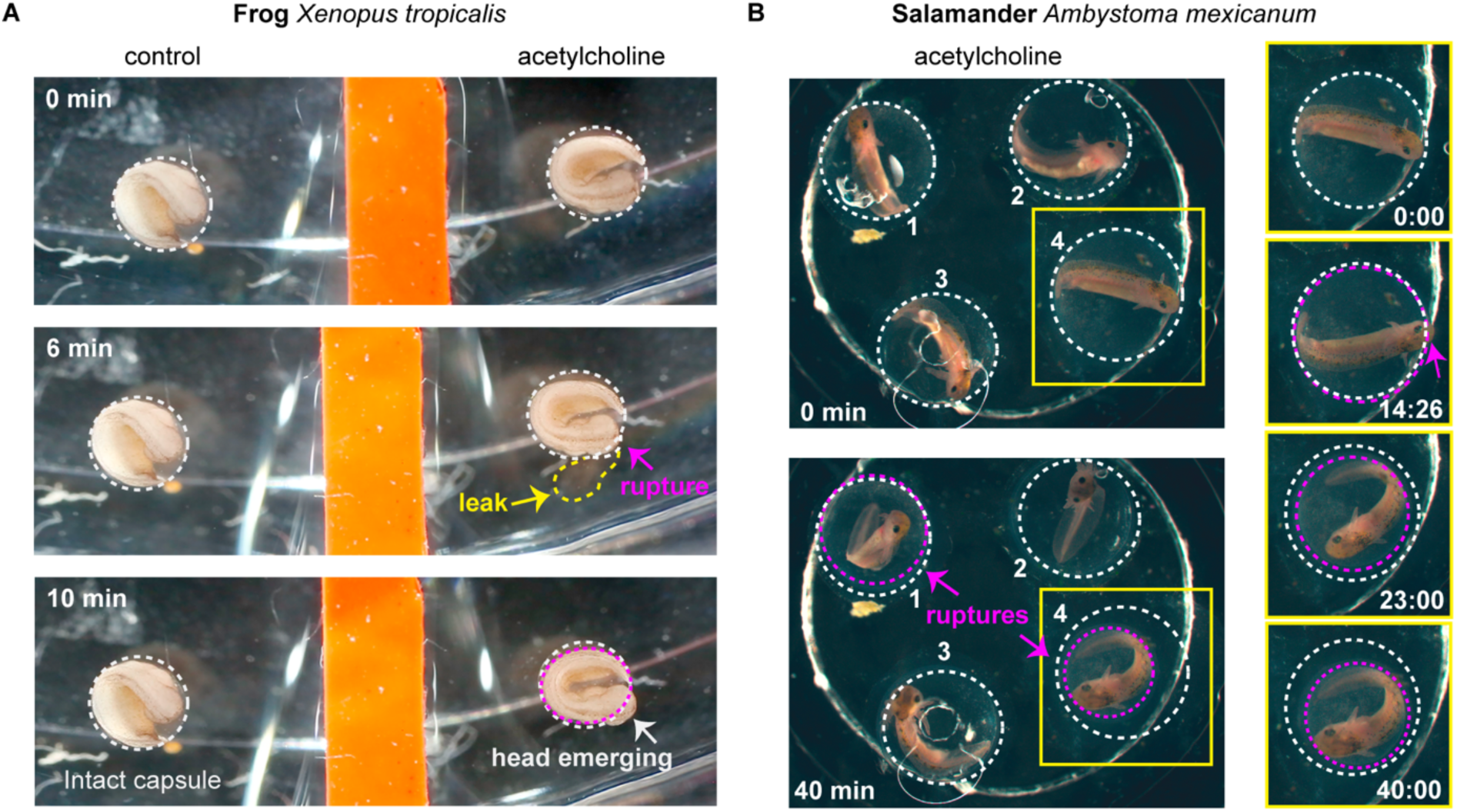
Acetylcholine triggers hatching enzyme release in distantly related amphibians. Acetylcholine induced egg membrane rupture in (A) the western clawed frog, *Xenopus tropicalis* and (B) the axolotl (salamander) *Ambystoma mexicanum.* Dashed lines indicate egg membrane positions (white: initial, magenta: shrinking ruptured membranes). Yellow arrow indicates perivitelline fluid leak, magenta arrows indicate rupture sites. Times from onset of exposure are indicated. (B) In axolotl, ruptures occurred in embryos 1 and 4; yellow boxes (right) show detailed timeline for embryo 4. Despite membrane ruptures, the jelly capsules surrounding membranes retained their integrity, containing the perivitelline fluid. No hatching occurred during the experiment.

## MOVIE CAPTIONS

**Movie S1**. Neurotransmitter screening trial showing acetylcholine-exposed *A. callidryas* embryo and paired sibling control. Time is from the start of acclimation and playback is accelerated; video starts when experimental solutions are in place. In acetylcholine, membrane rupture is evident as soon as the embryo moves and perivitelline fluid leaks (arrow) from where the embryo’s snout was initially positioned. The capsule collapses and embryo remains within, showing no hatching-specific behavior. The control egg remains intact.

**Movie S2**. Cholinergic receptor agonist trial showing carbachol (muscarinic agonist) exposed *A. callidryas* embryo and paired sibling control. Time is from the start of acclimation and playback is accelerated; video starts when experimental solutions are in place. In carbachol, membrane rupture is evident as soon as the embryo moves and perivitelline fluid leaks (arrow) from where the embryo’s snout was initially positioned. There is no hatching-specific behavior, and the embryo is removed after membrane collapse to avoid drug overexposure, then video ends. The control egg remains intact until the end of the trial (10 min, not shown).

**Movie S3**. Cholinergic receptor antagonist trials showing embryos’ response to: (1) a mechanosensory simulated predation cue (egg jiggling) and (2) a hypoxia cue, after 10 min exposure to either the muscarinic receptor antagonist (atropine) or nicotinic receptor antagonist (hexamethonium), each with a paired sibling control. Atropine and hexamethonium videos play concurrently; hexamethonium videos play continuously and atropine videos are spliced, removing periods without behavior. For the mechanosensory cue, time is from the onset of acclimation (3 min) before antagonist treatment. For the hypoxia cue, time is shown from onset of second acclimation (3 min) before flooding. For both, videos begin when cue begins. Playback speed is accelerated in all cases. Control and hexamethonium-exposed embryos perform hatching-specific behavior, rupture membranes, and exit normally. Atropine-exposed embryos repeatedly perform hatching-specific behavior, but capsules remain intact and embryos cannot exit.

**Movie S4.** Effects of serotonin and serotonin receptor 3 antagonist (MDL 72222) exposure (10 min) on embryos’ hatching response to hypoxia. Serotonin and MDL 72222 trials play concurrently. Experimental and paired sibling control embryos were submerged in hypoxic water for 30 min. Time is from the onset of cue and playback speed is accelerated. Antagonist video is spliced to shorten it after control hatches, removing a period when the experimental embryo did not move. Serotonin exposure accelerates the hatching response to hypoxia; MDL 72222 exposure delays or ablates it.

## STAR★METHODS

### KEY RESOURCES TABLE

| REAGENT or RESOURCE | SOURCE | IDENTIFIER |
|---|---|---|
| Antibodies |  |  |
| Rabbit anti-AcHE | DHSB | RRID: AB_3675360 |
| Rabbit anti-5HT | Sigma-Aldrich | Sigma-Aldrich: S5545 |
| Mouse anti-SV2 | DSHB | DSHB Cat# SV2, RRID: AB_2315387 |
| Mouse anti-Zn-12 | DSHB | DSHB Cat# zn-12, RRID:AB_531908 |
| Mouse anti-ACh | Abcam | Abcam: ab34935 |
| Phospho-S6 (Ser235, Ser236) Monoclonal Antibody (cupk43k), eBioscience™ | ThermoFisher Scientific | AB_2572910 |
| Goat anti-mouse Alexa fluor 555 | Invitrogen | Invitrogen: A-21422 |
| Goat anti-rabbit Alexa fluor 488 | Invitrogen | Invitrogen: A-11008 |
| Goat anti-rabbit Cy3 | Abcam | Abcam: ab6939 |
| Biological samples |  |  |
| *Agalychnis callidryas* eggs (wild type, field collected) | Experimental Pond, Gamboa, Panama | NA |
| *Agalychnis callidryas* eggs (wild type, captive bred) | from Sally Seraphin, Trinity College | NA |
| *Xenopus tropicalis* eggs (wild type) | Marine Biological Laboratory; https://www.xenopusnxr.org/xenbase/nxr/jsp/orderingAndPricing/orderInfo.jsp | Wild type embryos |
| *Ambystoma mexicanum* eggs (wild type) | Ambystoma Genetic Stock Center | Cat# AGSC_100E |
| *Danio rerio* eggs (wild type) | Zebrafish Facility, MCB, Harvard University | Wild type embryos |
| Chemicals, peptides, and recombinant proteins |  |  |
| Acetylcholine chloride, ACh | Sigma-Aldrich | A6625_Sigma |
| Serotonin (5-hydroxytryptamine hydrochloride), 5-HT | Sigma-Aldrich | Cat# H9523 |
| Noradrenaline bitartrate, NO | Tocris | Cat# 5169 |
| Dopamine hydrochloride, DO | Sigma-Aldrich | H8502 |
| Carbachol | Tocris | Cat# 2810 |
| Nicotine (nicotine ditartrate) | Tocris | Cat# 3546 |
| Atropine | Sigma-Aldrich | Sigma-Aldrich: A0257 |
| Hexamethonium chloride | Sigma-Aldrich | Sigma-Aldrich: H2138 |
| MDL 72222 (tropanyl 3,5-dichlorobenzoate), 5-HT3R antagonist | Tocris | Cat# 0640 |
| Dimethyl sulfoxide, DMSO | Sigma-Aldrich | Sigma-Aldrich: 472301 |
| MS-222 (ethyl 3-aminobenzoate methanesulphonate) | Sigma-Aldrich | Sigma-Aldrich: A5040 |
| Paraformaldehyde, PFA | Electron Microscopy Sciences | Cat# 157-4 |
| Phosphate-buffered saline, PBS | Sigma-Aldrich | Sigma-Aldrich: 6506-OP |
| Triton X-100, PBST | Sigma-Aldrich | Sigma-Aldrich: X100 |
| Normal goat serum, NGS 10% | Thermo Fisher Scientific | Cat# 500627 |
| Sodium Azide | Sigma-Aldrich | Sigma-Aldrich: S2002 |
| DAPI (4′,6-diamidino-2-phenylindole) | Sigma-Aldrich | Sigma-Aldrich: MBD0015 |
| Vectashield antifade mounting medium | Vector Laboratories | H-1200 |
| Vectashield antifade mounting medium with DAPI | Vector Laboratories | H-1200-10 |
| Glycerol | Sigma-Aldrich | Sigma-Aldrich:G5516 |
| Tissue-Tek® O.C.T. Compound | Electron Microscopy Sciences | Cat# 62550-12 |
| Gluteraldehyde-Paraformaldehyde | Electron Microscopy Sciences | Cat# 15720 |
| Critical commercial assays |  |  |
| RNAscope probe- Acall-*HE*-C1 *Agalychnis callidryas* hatching enzyme (AcHE) mRNA | Advanced Cell Diagnostics | Cat# 1216701-C1 |
| RNAscope probe- Acall-CHRM1-C2 *Agalychnis callidryas* muscarinic receptor 1 (mAChR1) mRNA | Advanced Cell Diagnostics | Cat# 1216681-C2 |
| RNAscope probe- Acall-CHRM2-C3 *Agalychnis callidryas* muscarinic receptor 2 (mAChR2) mRNA | Advanced Cell Diagnostics | Cat# 1216691-C3 |
| RNAscope probe- Xt-*CHRM1*-C2 *Xenopus tropicalis* muscarinic receptor 1 (mAChR1) mRNA | Advanced Cell Diagnostics | Cat#1573041-C2 |
| RNAscope probe- Am-CHRM1-C2 *Ambystoma mexicanum* muscarinic receptor 1 (mAChR1) mRNA | Advanced Cell Diagnostics | Cat# 1573261-C2 |
| RNAscope probe- Dr-CHRM1-C2- *Danio rerio* muscarinic receptor 1 (mAChR1) mRNA | Advanced Cell Diagnostics | Cat# 1565181-C2 |
| TSA-Cy3 | Perkin Elmer | Cat# NEL741001KT |
| TSA-Cy5 | Perkin Elmer | Cat# NEL741001KT |
| TSA-Cy5 | Perkin Elmer | Cat# NEL741001KT |
| RNAscope Multiplex Kit V2 | Advanced Cell Diagnostics | Cat# 323100 |
| Deposited data |  |  |
| Raw and analyzed data | This paper | BioProject: ID PRJNA1541026, Accession numbers SRR41071719 (RNAseq of *A.callidryas* gills), SRR41071720 (RNAseq of *A.callidryas* brain), SRR41071717 (RNAseq of *A.callidryas* tail skin), SRR41071721 (RNAseq of *A.callidryas* HGCs), SRR41071718 (RNAseq of *A.callidryas* yolk skin), SRR41071716 (RNAseq of *A.callidryas* lungs). |
| Experimental models: Organisms/strains |  |  |
| *Agalychnis callidryas* eggs (wild type, field collected) | Experimental Pond, Gamboa, Panama | NA |
| *Xenopus tropicalis* eggs (wild type) | Marine Biological Laboratory; https://www.xenopusnxr.org/xenbase/nxr/jsp/orderingAndPricing/orderInfo.jsp | Wild type embryos |
| *Ambystoma mexicanum* eggs (wild type) | Ambystoma Genetic Stock Center | Cat. No. AGSC_100E |
| *Danio rerio* eggs (wild type) | Zebrafish Facility, MCB, Harvard University | Wild type embryos |
| Software and algorithms |  |  |
| GraphPad Prism | GraphPad software | https://www.graphpad.com/ |
| Trinity v.2.4.0 | Grabherr et al.<sup>57</sup> |  |
| Kallisto | Bray et al.<sup>59</sup> |  |
| Diamond v.2.0.9 | Buchfink et al.<sup>58</sup> |  |
| FIJI ImageJ2 | Schneider et al.<sup>61</sup> | https://imagej.nih.gov/ij/ |

#### Red-eyed treefrogs

##### Egg collection and animal care

We collected *Agalychnis callidryas* egg clutches from the Experimental Pond (9.120894 N, 79.704015 W) at the Smithsonian Tropical Research Institute (STRI) in Gamboa, Panama, in the 2021–2023 rainy seasons. We transported clutches to a nearby ambient laboratory where we removed any unfertilized eggs, mounted each clutch on a plastic card and placed it in an individual cup over dechlorinated tap water. We maintained clutches in egg humidors (plastic boxes with partially screened lids) where they were misted with rainwater multiple times daily, either manually (2021) or automatically, using a MistKing system (2022, 2023; https://junglehobbies.com/). We adjusted misting frequency as needed to maintain optimal egg hydration, based on weather, and removed excess water from the cups twice daily. All embryos used in experiments were healthy and developing normally. At the end of the trial, we manually decapsulated any unhatched embryos. We euthanized a subset of individuals using an overdose of MS-222 (1 g/L, buffered to pH 7.0–7.5) and preserved them for histological analysis. We reared all other tested individuals for two weeks in plastic cups with dechlorinated water, keeping each treatment separate to monitor for normal development, and provided Sera Micron tadpole food ad libitum once tadpoles began feeding. Then we released them back into the Experimental Pond. Additional, untested individuals from these clutches were returned to the pond immediately after hatching. All research in Panama was approved by the STRI ACUC (protocol 2020-0318-2023) and permitted by the Panamanian Ministry of the Environment (research: ARB-044-2021, specimen export: 01-ARG-090-2021, PA01ARB0146-2022).

#### Behavioral pharmacology trial setup and drug exposure

We used pairs of stage-matched experimental and control embryos from the same clutch for each trial. Immediately before each trial, we carefully removed each egg from the clutch using curved forceps, de-jellied it, placed it in either a hexagonal weigh boat or well of a 24-well cell culture plate, and let it acclimate for 2–3 minutes. Pairs of weigh boats were glued onto shared base plates for stability. After acclimation, we added ∼750 µl of control (usually water) or experimental solution, using a fine tip 1.5 ml transfer pipet, to submerge the bottom half of the egg, leaving the top exposed to air. We determined the exposure time (10 min in all cases) and drug concentration for each neurotransmitter, agonist, and antagonist based on pilot trials. Our aim was to allow enough drug to enter the egg capsule and reach the embryo’s epithelial cells (hatching gland cells on head or neuroepithelial cells on gills) to get a clear response, while avoiding overexposure that might lead to unintended effects on the embryo’s physiology or behavior. We left unhatched eggs in solutions for 10 minutes, then removed the solution and ended the trial or proceeded with the following experimental step (hatching cue or second drug solution) as applicable. If an individual hatched before 10 minutes, we removed it to prevent further drug exposure.

#### Video recording

We video-recorded behavioral pharmacology trials for analysis, using a Canon EOS Mark III or Mark IV camera and a Canon MP-E 65 mm or 100 mm lens, illuminating the test arena (well plate, weigh boats, or aquarium) from both sides using two LED lights. For uncued trials, we began recording during acclimation to the drug-exposure arena and continued through the end of exposure. For cued trials, we started recording after drug treatment, during the second acclimation, and continued through the end of cue exposure.

#### Uncued behavioral pharmacology experiments

##### Screen for neurotransmitter and receptor agonist effects on hatching enzyme release

We exposed experimental embryos to four neurotransmitters – acetylcholine (1 mM), serotonin (500 μm), noradrenaline (1 mM), and dopamine (1 mM) – to test their potential role in hatching enzyme release, monitoring eggs for capsule rupture. We selected neurotransmitters to test based on acetylcholine’s common role in glandular secretion^23^, norepinephrine’s role in anuran mucus and granular glands^24^, dopamine’s hypothesized role in fish hatching^25^, and serotonin’s role in hypoxia-cued behavior in fishes^26^. We used embryos from 21 clutches and conducted trials starting at stage 28 (age 4 days, Warkentin et al.^50^). Stage 28 embryos hatch in response to both hypoxia and mechanosensory cues^18, 20^ but are less vibration sensitive than later in development^22, 40, 51, 52^, facilitating experimental manipulation. Sample sizes were 31 pairs of embryos tested with acetylcholine, 13 pairs with serotonin, and 17 pairs with noradrenaline (all at age 4 days), plus 10 pairs with dopamine at early 5 days. Following positive results with acetylcholine, we performed trials exposing 4-day-old embryos to muscarinic and nicotinic cholinergic receptor agonists, specifically carbachol (5mM, 25 pairs) and nicotine (500 μm, 16 pairs). A subset (N=6) of nicotine-exposed pairs were exposed to a mechanosensory hatching cue (see below) following trials to assess hatching competence, and hatched tadpoles’ swimming was observed to verify normal mobility. In carbachol trials, after membrane rupture and capsule collapse embryos were removed to avoid overexposure to the drug.

#### Cued behavioral pharmacology experiments

To assess the effects of specific neurotransmitters, agonists, and antagonists on components of the cued hatching process, we followed drug exposures with two natural hatching cues, hypoxia and mechanosensory stimulation.

##### Hypoxia cue

Following Warkentin et al.^18^, we flooded eggs with hypoxic water (ambient temperature, ca. 15% air saturated) to examine hypoxia-cued hatching. To stabilize eggs for video-recording and eliminate motion cues during trials, we used a custom-made “egg bridge” constructed from a glass arch mounted on a glass base, with two loops of nylon monofilament stretched around the parallel vertical columns of the arch. After drug exposure, we placed test eggs on the bottom loop, then slid the top loop down to hold them in place, ensuring that the eggs were not in contact. The bridge-base was embedded in ThermoMorph plastic weighted with glass marbles, custom fit to the bottom of a small aquarium (10.3×4.9×8 cm in width, depth, height). This set-up prevented egg movement during flooding and provided complete egg-surface exposure and excellent visibility. We used stage 26–28 embryos for these experiments (Warkentin et al.^50^; late 3 to 4 days old). These stages consistently hatch in response to hypoxia^50^ but are unresponsive or minimally responsive to mechanosensory cues^20^, so egg-transfer to the bridge does not induce hatching. We allowed embryos 3 min of acclimation, then poured in hypoxic water to completely submerge them. Under these conditions, hatched tadpoles rapidly surface to fill their lungs. To avoid disturbance of unhatched embryos, we removed tadpoles as soon as they surfaced, sucking them into a wide-mouthed transfer pipette. After 30 min of submergence, we removed unhatched eggs from the bridge and manually decapsulated embryos. We immediately transferred hatched and decapsulated animals to normoxic water.

##### Mechanosensory simulated predation cue

Following Warkentin et al.^18^, we used a standardized egg-jiggling protocol, simulating a predator attack. We conducted all mechanosensory trials in weigh boats. After drug exposure, we removed the solutions and left embryos undisturbed for 3 min, then jiggled eggs intermittently, in bouts of a few seconds, using moistened, blunt-ended, metal probes and alternating between control and experimental embryos. We stopped stimulation when an embryo started shaking, a characteristic hatching-specific behavior^12, 22^, and resumed it if the embryo ceased shaking without hatching. After 10 min, we ceased stimulation and manually decapsulated unhatched embryos.

#### Effect of cholinergic receptor blocking on hatching enzyme release

We exposed embryos to specific antagonists of muscarinic receptors (atropine, 100 μm) or nicotinic receptors (hexamethonium chloride, 500 μm), then cued them to hatch, monitoring for hatching-specific behavior and membrane rupture. We used two different hatching cues to assess if both sensory modalities activate the same enzyme-release mechanism. For hypoxia trials, we used 4-day-old embryos (21 pairs for atropine, 9 for hexamethonium) to avoid hatching during transfer to the egg-bridge. For mechanosensory trials, we used 5-day-old embryos which are highly responsive to mechanosensory stimuli (25 pairs for atropine, 13 for hexamethonium).

#### Effect of serotonergic signaling on the hatching response to hypoxia

We modified Regan et al.^53^ amphibious fish protocols for use with terrestrial anuran embryos. We pre-exposed embryos to serotonin (500 μm, 23 pairs) or the serotonin receptor 3 antagonist MDL 72222 (tropanyl 3,5-dichlorobenzoate, 500 μm, 21 pairs), then tested their response to hypoxia. We used DMSO (final concentration ≤ 0.5%) as a vehicle for MDL 72222, so paired controls included this vehicle. In addition, we did standard water controls in a subset of trials, using trios of stage-matched siblings, and found no effect of the vehicle (data not shown).

#### Effect of serotonin receptor 3 antagonist on mechanosensing and hatching enzyme release

To test if the serotonin receptor 3 antagonist (MDL 72222) affects the hatching response to a non-hypoxia cue, via cue sensing and/or hatching enzyme release mechanisms, we pre-exposed embryos to MDL 72222 (500 μm), then exposed them to acetylcholine (1 mM, 8 pairs, 4 days old) or egg-jiggling (6 pairs, 5 days old).

#### Video analysis

From video recordings, we quantified three key elements of the hatching process^12^. As an indicator of the decision to hatch, we recorded the occurrence (or not) of a hatching-specific shaking and pushing behavior that embryos commonly perform just before membrane rupture, with their hatching gland cells positioned against the vitelline membrane^12, 19, 22^. As an indicator of hatching enzyme release, we recorded the occurrence of membrane rupture, identifying it by the leakage of perivitelline fluid or protrusion of part of the embryo’s head outside the membrane. We also recorded the timing of the initial membrane rupture and noted any additional ruptures. Finally, we recorded if and when the embryo exited from its egg capsule, successfully completing the hatching process^12, 22^. In the neurotransmitter screening, for embryos that exited the capsule, we noted if exit immediately followed rupture, through the initial rupture site, or if the embryo moved away from the initial rupture site, allowing fluid leakage and capsule collapse, and then remained within the capsule for some time before exiting through a subsequent rupture; we recorded the latter as a delayed exit.

#### Specimens for visualization of hatching enzyme reserves

We euthanized embryos after three treatments to visualize the effects of acetylcholine and a muscarinic receptor antagonist (atropine) on hatching enzyme reserves^19^. Embryos were either manually decapsulated and then exposed to acetylcholine for 10 min, exposed to atropine for 10 min in *ovo*, or exposed to atropine in *ovo* followed by 5 min of egg-jiggling plus 10 min of acetylcholine exposure.

#### Tissue preparation for immunohistochemistry

Whole embryos or dissected gill tissue (internal plus attached external gills) were fixed in 4% paraformaldehyde prepared in 1x phosphate buffered saline (PBS) at 4°C for 12–24 h, then rinsed for 12–24 h in PBS at 4 °C. Samples were used immediately or stored in PBS with 0.1% sodium azide at 4 °C, then rinsed in PBS for 24h at 4 °C before use. To prepare sections, we cryoprotected whole embryos in 30% sucrose for 24 h, then froze them in OCT. We trimmed blocks and sectioned specimens at 18–50 µm using Leica CM1860 or CM3050S cryostats. We rinsed sections on slides in PBS before staining.

#### Immunohistochemistry

For immunostaining, we permeabilized samples at 4 °C in 0.5% Triton X-100 in PBS (PBST) with 1% normal goat serum (NGS) for 48–72 h, then placed them in a blocking solution of PBST + 5 % NGS for 2 h on a shaker at room temperature (RT). We incubated samples in primary antibody with PBST for 48–72h at 4 °C, rinsed them with PBST, then applied the secondary antibody in the dark on a shaker for 4h at RT or overnight at 4 °C, and rinsed with PBS. We counter-stained whole embryos in DAPI for 30 min in the dark on a shaker, then rinsed with PBS again. We placed whole embryos in 60% glycerol with PBS overnight at 4 °C, then in 90% glycerol at 4 °C until imaging. We mounted gills and sections on glass slides in Vectashield antifade mounting medium, with or without DAPI, and stored them at 4 °C until imaging. To mark the hatching enzyme reserves, we used rabbit anti-AcHE (1:200 in whole embryos or 1:500 for cryosections)^19^ and visualized it with goat anti-rabbit Alexa fluor 488 (1:500). To mark nerves on the snout, we used mouse anti-Zn-12 (1:50) to mark nerve cell surfaces and mouse anti-SV2 to mark synaptic vesicles; to identify cholinergic nerves near the hatching gland cells, we used mouse anti-ACh to mark acetylcholine. We visualized these with goat anti-mouse Alexa fluor 555 and 594 (both 1:500). We examined gill innervation using mouse anti-Zn-12 and identified serotonergic neuroepithelial cells using rabbit anti-5-HT (1:500) and mouse anti-SV2^54, 55^. We visualized markers with goat-anti-rabbit Alexa fluor 488, goat anti-rabbit Cy3, and goat anti-mouse Alexa fluor 555. We imaged gills, sectioned tissue, and whole-mount embryo samples with FV1000 or FV3000 (Olympus Life Sciences, Smithsonian Tropical Research Institute), LSM 900 Confocal Microscope (Zeiss, Harvard Center for Biological Imaging) confocal microscopes. We processed the images using Fiji (NIH).

#### Transmission electron microscopy

We fixed whole, 5-day-old embryos collected in Gamboa, Panama, in a mixture of 2% formaldehyde, 2.5% glutaraldehyde in 0.1 M sodium cacodylate buffer, pH 7.4. We processed the fixed specimens at the Electron Microscopy Lab (Harvard Medical School), using the following protocol: Samples were washed with 0.1 M sodium cacodylate buffer and post-fixed with 1% osmium tetroxide, 1.5% potassium ferrocyanide in H_2_O for 2 h. The samples were then washed in a maleate buffer and post-fixed in 1% uranyl acetate in maleate buffer for 1 h, rinsed in double-distilled H_2_0, and dehydrated through a series of ethanol (50%, 70%, 95% and twice at 100%) for 15 min per solution. Dehydrated tissues were placed in propylene oxide for 5 minutes before infiltration with Epon mixed 1:1 with propylene oxide overnight at 4 °C, then polymerized in a 60 °C oven in Epon resin for 48 h. They were then sectioned into 80-nm-thin sections and imaged on a JEOL 1200EX Transmission Electron Microscope. Tissues from three different embryos were analyzed.

#### Transcriptomics

We generated tissue-specific de novo transcriptomes for *A. callidryas* embryos. To obtain enough hatching gland cells, we dissected and pooled tissues from 400 embryos (5 days old, ≥ stage 30, from 10 clutches) in Panama. Tissues included: (1) a patch of snout skin where hatching gland cells are densely concentrated, medially above the mouth and between (not including) the naris and supraorbital neuromast lines (Figure 3A), (2) most of the tail skin, including neuromasts and lacking hatching gland cells, (3) most of the belly skin surrounding the yolk sac, without hatching gland cells or neuromasts, (4) the entire brain, (5) the gills, including internal and attached external gills, (6) the lungs. We placed tissues in RNAlater at 4 °C for 24 h, then at –20 °C until exportation on dry ice.

RNA extraction, library preparation, and RNA sequencing were performed by Genewiz (Azenta) using a HiSeq (2×150 bp) platform. The RNA sequencing libraries were prepared using the NEBNext Ultra II RNA Library Prep Kit for Illumina using manufacturer’s instructions (New England Biolabs, Ipswich, MA, USA). Briefly, mRNAs were initially enriched with Oligod(T) beads. Enriched mRNAs were fragmented for 15 minutes at 94 °C. First strand and second strand cDNA were subsequently synthesized. cDNA fragments were end repaired and adenylated at 3’ends, and universal adapters were ligated to cDNA fragments, followed by index addition and library enrichment by PCR with limited cycles. The sequencing library was validated on the Agilent TapeStation (Agilent Technologies, Palo Alto, CA, USA) and quantified by using Qubit 2.0 Fluorometer (ThermoFisher Scientific, Waltham, MA, USA) as well as by quantitative PCR (KAPA Biosystems, Wilmington, MA, USA).

The sequencing libraries were multiplexed and clustered on 1 lane of a flowcell. After clustering, the flowcell was loaded on the Illumina HiSeq 4000 instrument according to manufacturer’s instructions. The samples were sequenced using a 2×150 Pair-End (PE) configuration. Image analysis and base calling were conducted by the HiSeq Control Software (HCS). Raw sequence data (.bcl files) generated from Illumina HiSeq was converted into fastq files and de-multiplexed using Illumina’s bcl2fastq 2.17 software. One mismatch was allowed for index sequence identification.

We trimmed adaptors using TrimGalore^56^ and assembled reference transcriptomes de novo using Trinity^57^. We identified open reading frames using TransDecoder^58^ and used Kallisto^59^, with our novel transcriptome assembly as a reference, to pseudo-align reads and estimate transcript abundance. Annotation was performed using Diamond^58^. Sequencing data generated from this study are available at the NCBI Sequence Read Archive under BioProject ID PRJNA1541026, Accession numbers SRR41071719 (RNAseq of *A.callidryas* gills), SRR41071720 (RNAseq of *A.callidryas* brain), SRR41071717 (RNAseq of *A.callidryas* tail skin), SRR41071721 (RNAseq of *A.callidryas* HGCs), SRR41071718 (RNAseq of *A.callidryas* yolk skin), SRR41071716 (RNAseq of *A.callidryas* lungs).

#### Comparative studies

To assess the potential phylogenetic generality of cholinergic signaling for hatching enzyme release in anamniotic vertebrates, we worked with aquatic embryos of a distantly related model frog, salamander, and fish (*Xenopus tropicalis, Ambystoma mexicanum*, and *Dario rerio*) at Harvard University (IACUC protocol 18-12-343). We tested each species for the effect of acetylcholine on hatching enzyme release. We adapted the methods developed for *A. callidryas* (above) as needed to account for differences in egg habitat and structure, developmental timing, and hatching process. Specifically, we incubated dejellied *X. tropicalis* eggs in Petri dishes with 0.1% MMR at 18–23 °C. We reared *A. mexic*anum eggs in 20% Holtfreter’s solution in plastic containers at 23 °C. We incubated *D. rerio* eggs in Petri dishes with E3 medium (5 mM NaCl, 0.17 mM KCl, 0.33 mM CaCl_2_, 0.33 mM MgSO_4_, and 120 μl/L 0.1% methylene blue)^60^ at 23 °C. We checked all eggs daily for normal development, removing any unfertilized, abnormally developing, or damaged eggs. Acetylcholine (1 mM) exposures were as follows: *X. tropicalis* (9 pairs) for 10 min, *A. mexicanum* (12 pairs) for up to 45 min, and *D. rerio* (6 pairs) for up to 40 min. We recorded the trials using a Canon EOS Mark IV camera with a Canon MP-E 65 mm macro lens (*X. tropicalis*) or an AxioZoom V.16 microscope (*A. mexicanum* and *D. rerio*).

#### Fluorescence in situ hybridization (RNAscope)

To localize muscarinic acetylcholine receptors in *A. callidryas*, *X. tropicalis, A. mexicanum* and D*. rerio*, we used RNAscope with anti-HE, anti-mAChR1, and anti-mAChR2 probes. Captive-bred *A. callidryas X. tropicalis, A. mexicanum*, embryos were euthanized and decapsulated, then immediately frozen in OCT and stored at -80 °C until use. For zebrafish, embryos were manually dechorionated and fixed in 4% PFA for 2 hours at room temperature to avoid the yolks dissolving in OCT. We trimmed blocks and sectioned at 18–50 µm using a Leica CM3050S cryostat. We used probes designed by ACD Inc., based on hatching enzyme, mAChr1, and mAChR2 sequences from our *A. callidryas* transcriptome, or based on published sequences for *X. tropicalis, A. mexicanum*, and *D. rerio* mAChR1; hatching enzyme probe from *A. callidryas* cross-reacts with the other three species. We followed the manufacturer-recommended protocol for fresh frozen tissues. We used the pretreat 3 for 30 min and the included fluorescent probes TSA-Cy3, TSA-Cy5, and TSA-FITC.We mounted samples in Vectashield with DAPI (Vector Labs #H-1200-10) and imaged them with an LSM 900 Confocal Microscope (Zeiss), or AxioZoom v16 microscope (Harvard Center for Biological Imaging). Images were processed using FIJI (NIH).

#### Quantification and statistical analysis

We used GraphPad Prism v10.02 for Mac (www.graphpad.com) for statistical analysis. We used a Kruskal-Wallis test followed by Dunn’s multiple comparisons to compare fluorescence levels in hatching gland cells exposed to atropine, acetylcholine, or both (Figure 2D-E). We used Wilcoxon matched-pairs signed rank tests for non-parametric data to compare membrane rupture time between control and treatment embryos in serotonin (Figure 3F) and Fisher’s exact test to compare hatching, membrane rupture or observed hatching-specific behavior proportions between in serotonergic and cholinergic antagonists and controls (Figure 3G and 4A).

